# The Left Insula Bridges Cognition, Emotion, and Brain Structure: A Multilayer Network Analysis of the Human Connectome Project-Young Adult

**DOI:** 10.1101/2024.10.30.621202

**Authors:** Lyndon Firman-Sadler, Mervyn Singh, Juan F. Domínguez D, Ivan L. Simpson-Kent, Kasren Caeyenberghs

## Abstract

Multilayer network analyses allow for the exploration of complex relationships across different modalities. Specifically, this study employed a novel method that integrates psychometric networks with structural covariance networks to explore the relationships between cognition, emotion and the brain. Psychological (NIH Toolbox Cognition Battery and NIH Toolbox Emotion Battery) and anatomical MRI (cortical volume) data were extracted from the Human Connectome Project Young Adult dataset (*n* = 1109). Partial correlation networks with graphical lasso regularisation and extended Bayesian information criterion tuning were used to model a psychometric bi-layer network consisting of seven cognitive nodes and four emotion nodes, as well as a neuro-psychometric tri-layer network consisting of these same nodes in addition to 24 brain nodes from the central executive and salience networks. Bridge strength centrality was used to identify nodes that bridged between layers. For the bi-layer network, it was found that stress was the only bridge node. For the tri-layer network, six bridge nodes were identified, with the left insula emerging as the most central. These findings demonstrate the utility of multilayer networks in integrating psychological and neurobiological data for the potential identification of targets to improve psychological wellbeing.

## 1. Introduction

The symptomatology of many clinical conditions includes changes to cognition and emotion. For example, in neurological conditions such as following from a stroke or traumatic brain injury, individuals suffer from memory impairments, as well as low mood and anxiety (Konrad et al., 2011; Terrill et al., 2018). Similarly, psychological conditions such as Major Depressive Disorder, Generalised Anxiety Disorder, and schizophrenia involve both cognitive and emotional symptoms (American Psychiatric Association [APA], 2013; Diener et al., 2012). The relationships between cognition and emotion in these conditions are often complex and bi-directional. For example, after a stroke, individuals may have cognitive impairments, such as limited attention and working memory capacity that cause difficulties with daily tasks, leading to negative emotions, such as sadness and fear (Terrill et al., 2018). Conversely, these negative emotions may in turn exacerbate attention difficulties, as is often seen in Major Depressive Disorder (APA, 2013; Wang et al., 2020).

Network science has demonstrated utility in understanding how psychological constructs such as cognition and mental health are subserved by complex relationships between components (Ferguson, 2021; Giles et al., 2022; Kan et al., 2019; Kenny et al., 2021). Psychometric networks often use partial correlation networks (Epskamp & Fried, 2018) to analyse the conditional dependencies of psychological variables such as cognitive abilities (e.g., working memory, attention, etc.) or mental health symptoms (e.g., anxiety, depression, etc.). The variables (nodes) in these networks are connected to other nodes by edges, which represent the statistical relationships between them (i.e., partial correlations). The importance of nodes in the network can then be quantified by centrality metrics, such as strength centrality, which sums the absolute weight of the edges from each node to all other nodes in the network (Epskamp & Fried, 2018). Bridge strength centrality specifically identifies nodes that are central to connecting different constructs, such as between comorbid conditions (Jones et al., 2021).

Psychometric networks have been used to better understand the relationships between the cognitive abilities involved in intelligence (Kan et al., 2019; Schmank et al., 2019), the relationships between symptoms within a number of mental health conditions, such as depression (Kenny et al., 2021), eating disorders (Giles et al., 2022), borderline personality disorder (De Paoli et al., 2020), and obsessive-compulsive disorder (Giles et al.). The approach has also been used in neurodegenerative conditions such as Alzheimer’s Disease (Ferguson, 2021) where, for example, researchers identified processing speed as the most central node in the network for those with Alzheimer’s Disease, and episodic memory as the most central node for controls (Ferguson, 2021). These findings suggest that psychometric networks can both identify cognitive abilities important to normal functioning and detect how these cognitive functions are altered in disease.

Psychometric networks are therefore a suitable framework to look at the complex and dynamic relationships between cognition and emotion. However, to truly understand the link between cognition and emotion, it is also necessary to identify the brain regions that are involved in both. It has been argued by some researchers that cognitive and emotional processes arise from often overlapping neurophysiological functions, and therefore, there are likely brain regions responsible for both cognitive abilities and emotions (Barrett, 2016; Kraljević et al., 2021; Pessoa, 2008). Brain regions within the central executive and salience networks have previously been found to be associated with a mixture of cognitive abilities and emotional states (Cromheeke & Mueller, 2014; Diener et al., 2012; Kraljević et al., 2021; Metzler-Baddeley et al., 2016). Furthermore, disruptions to these brain networks have been implicated in the cognitive or emotional symptoms of numerous psychopathological conditions (Menon, 2011). As neuroimaging studies typically focus on the neural corelates of either cognition or emotion, few focus on identifying brain regions involved in both (Diener et al., 2012; Kraljević et al., 2021). However, one such study by Kraljević et al. (2021) used multiple linear regression models with data from the Human Connectome Project and found that the superior frontal cortex is significantly associated with both cognition and emotion. This discovery highlights the potential for further exploration of the relationship between cognition, emotion, and the brain.

A small number of studies have begun to model networks with both psychometric and neuroimaging data (Bathelt et al., 2022; Hilland et al., 2020; Simpson-Kent et al., 2021; for overviews see Blanken et al., 2021; Vitevitch, 2025). One approach is to integrate psychometric networks with structural covariance networks in a multilayer network (Hilland et al., 2020; Simpson-Kent et al., 2021). For example, Simpson-Kent et al. (2021) used multilayer partial correlation networks with a range of cognitive and neuroimaging measures from a sample of 805 students (ages 5-18 years) with learning difficulties. Specifically, the authors looked at the relationships between cognitive abilities, cortical volume, and fractional anisotropy. Reading ability was found to be the most central node, suggesting that for those with learning difficulties, interventions that focus on improving reading may lead to improvements in other cognitive abilities. Additionally, the superior frontal gyrus was found to be a bridge node, which suggests this region may be a target for intervention through non-invasive brain stimulation or cognitive training tasks that activate this area (Simpson-Kent et al., 2021).

Although a small number of studies have begun to explore the integration of psychological and neuroscientfic data together within a unified network framework (Bathelt et al., 2022; Blanken et al., 2021; Hilland et al., 2020; Simpson-Kent et al., 2021), no study has yet explored the integration of a psychometric network consisting of both cognitive abilities and emotion, with a brain network. Such an exploration of the relationship between cognition and emotion allows for a greater understanding of the complex relationships often seen in clinical populations but also in non-clincial and sub-clinical contexts, potentially aiding in identifying targets for intervention. In addition, by considering the involvement of brain regions in this interplay, it may be possible to elucidate not only additional treatment targets but also establish a deeper understanding of the complex relationship between the brain and both cognition and emotion. Lastly, establishing a multilayer network framework of psychometric and brain structural relationships with a large sample from a non-clinical young adult population can provide a baseline for comparison in future studies involving clinical samples and other age groups.

Using a large sample of healthy young adults from the Human Connectome Project, the present study therefore aims to explore the relationships between general cognitive abilities (Diener et al., 2012; Konrad et al., 2011) and the negative emotions often implicated in psychopathology (Diener et al., 2012; Salsman et al., 2013; Terrill et al., 2018), as well as their associated brain regions. To achieve this aim, we will first analyse a bi-layer network of cognitive abilities and negative emotions to determine the nodes that are most central in bridging cognition and negative emotion. Secondly, we will analyse a tri-layer network, consisting of cognitive abilities, negative emotions, and cortical brain regions from the central executive and salience networks (Cromheeke & Mueller, 2014; Diener et al., 2012; Kraljević et al., 2021; Metzler-Baddeley et al., 2016), to determine the nodes that are most central in bridging cognition, emotion and the brain.

## 2. Materials and Methods

### 2.1. Participants

The present study utilised pre-existing data from the Human Connectome Project (http://www.humanconnectome.org), specifically the Young Adult cohort (HCP-YA; publicly available since February 2017). The data acquisition for HCP-YA was led by Washington University and the University of Minnesota (Van Essen et al., 2012). Processed data was accessed for the 1206 HCP-YA participants. However, only participants with complete data across all variables were included in the analysis. Ninety-three participants were excluded as they did not have anatomical MRI data. Four participants were excluded as they were missing psychological data. The remaining 1109 participants were aged between 22 and 35 years. There were 506 males (45.63%) and 603 females (54.37%). The participants included in HCP-YA were considered healthy, with no history of significant psychiatric or neurological conditions (Van Essen et al., 2012). Furthermore, the age range of participants was chosen to include adults during a period of relatively minimal neurological change, i.e., past the age of major neurodevelopmental changes and before the age of neurodegeneration (Van Essen et al., 2012). As this study only used pre-existing data, ethics exemption was granted through the Deakin University Human Research Ethics Committee.

### 2.2. Psychometric Measures

Cognition and emotion data in HCP-YA were acquired with the National Institute of Health (NIH) Toolbox for Assessment of Neurological and Behavioral Function® (NIH Toolbox; neuroscienceblueprint.nih.gov), a well-validated and reliable battery of measures (Salsman et al., 2013; Weintraub et al., 2013) that can be used with healthy and clinical populations. The NIH Toolbox can also be used to examine relationships between psychological functions and brain structure (Barch et al., 2013). The present study used data obtained with two of the NIH Toolbox’s modular batteries: the NIH Toolbox Cognition Battery and the NIH Toolbox Emotion Battery.

#### 2.2.1. Cognition

All seven of the cognitive tasks (Table 1) from the NIH Toolbox Cognition Battery (Weintraub et al., 2013) were included in analyses to capture a broad range of general cognitive abilities that vary in both healthy populations and in psychopathology (Al-Qazzaz et al., 2014; Cassetta et al., 2020; Diener et al., 2012). The tasks included the Picture Sequence Memory Test (episodic memory), Dimensional Change Card Sort Test (executive functioning), Flanker Task (attention), Oral Reading Recognition Test (reading), Picture Vocabulary Test (vocabulary), Pattern Comparison Processing Speed Test (processing speed), and List Sorting Working Memory Test (working memory). Scores for each variable were converted to *t*-scores, with higher scores indicating better performance.

**Table 1.** Cognitive variables as measured by the NIH Toolbox Cognition Battery.

| Cognitive Variable | NIH Task | Task Description | Scoring |
| --- | --- | --- | --- |
| Episodic memory<br>(The ability to acquire, store and retrieve new information.) | NIH Toolbox Picture Sequence Memory Test | Participants are required to recall a sequence of pictures in the same order as originally presented, over two trials. | Scoring is based on the number of pictures recalled in the correct sequence, with higher scores indicating better episodic memory. |
| Executive function<br>(The ability to plan, organise and monitor the execution of behaviours to achieve a goal.) | NIH Toolbox Dimensional Change Card Sort Test | Participants are required to match a picture to one of two pictures, according to either colour or shape. The dimension for matching alternates between trials. | Scoring is based on a combination of accuracy and reaction time, with greater scores indicating better executive control. |
| Attention<br>(The ability to selectively attend to the appropriate stimuli amidst an abundance of stimuli.) | NIH Toolbox Flanker Inhibitory Control and Attention Test | Participants are required to respond in the direction of a middle arrow, that is flanked by additional arrows that are pointing in either the congruent or incongruent direction of the middle arrow. | Scoring is based on a combination of accuracy and reaction time, with greater scores indicating better attention. |
| Reading<br>(The ability to accurately read and pronounce words.) | NIH Toolbox Oral Reading Recognition Test | Participants are required to read aloud the word on the screen. | Scoring is based on correctly reading and pronouncing the word, with greater scores indicating better reading ability. |
| Vocabulary<br>(The ability to recall the meaning of words based on previous learning.) | NIH Toolbox Picture Vocabulary Test | Participants hear a word and are required to select, from four images, the image that most closely represents the heard word. | Scoring is based on selecting the correct image, with greater scores indicating better vocabulary. |
| Processing speed<br>(The speed at which one is able to process and respond to information.) | NIH Toolbox Pattern Comparison Processing Speed Test | Participants are required to identify whether two images are identical. | Scoring is based on the number of items correctly answered in 90 seconds, with greater scores indicating faster processing speed. |
| Working memory<br>(The ability to temporarily hold information in memory so that it can be utilised.) | NIH Toolbox List Sorting Working Memory Test | Participants are first required to sequence either food or animals in order of size based on remembering an unordered sequence of images. A second series of trials involves remembering both food and animals. | Scoring is based on the number of correctly ordered items in both series, with greater scores indicating better working memory. |

#### 2.2.2. Emotion

Four negative emotion scales (Table 2) were selected from the NIH Toolbox Emotion Battery (Salsman et al., 2013), i.e., the Anger-Affect Survey (anger), Fear-Affect Survey (fear), Sadness Survey (sadness), and Perceived Stress Survey (stress). Scales for negative emotion were included because they are able to capture variations in mental health in non-clinical contexts and are able to translate to clinical contexts as representations of symptoms (Salsman et al., 2013). For example, items within the Sadness Survey have been used to assess depression and items within the Fear-Affect Survey have been used to assess anxiety (Salsman et al., 2013). Scores for each emotion variable were converted to *t*-scores, with higher scores indicating more frequent experiences of that emotion.

**Table 2.** Emotion variables as measured by the NIH Toolbox Emotion Battery.

| Emotion Variable | NIH Survey | Item Description | Scoring |
| --- | --- | --- | --- |
| Anger<br>(An affect involving hostility and cynicism, which often involves frustration at goal-directed behaviour being impeded.) | NIH Toolbox Anger-Affect Survey | Rated on a five-point Likert scale, each item begins with 'in the past 7 days...' followed by a prompt relating to experiences of anger, e.g., 'when I was frustrated, I let it show'. | Higher scores indicate more frequently experienced anger. |
| Fear<br>(An affect characterised by autonomic arousal in response to perceived threat, which is considered to be an experience of anxiety when the response is disproportionate to the threat.) | NIH Toolbox Fear-Affect Survey | Rated on a five-point Likert scale, each item begins with 'in the past 7 days...' followed by a prompt relating to experiences of fear or anxiety, e.g., 'I felt anxious'. | Higher scores indicate more frequently experienced fear/anxiety. |
| Sadness<br>(An affect characterised by low mood, as well as negative self-perceptions and pessimism, which can be indicators for depression.) | NIH Toolbox Sadness Survey | Rated on a five-point Likert scale, each item begins with 'in the past 7 days...' followed by a prompt relating to experiences of sadness, e.g., 'I felt worthless'. | Higher scores indicate more frequently experienced sadness. |
| Stress<br>(Stress is an individual's perception of being overwhelmed by life's demands, without sufficient resources to meet the demands or effectively cope with the pressure.) | NIH Toolbox Perceived Stress Survey | Rated on a five-point Likert scale, each item begins with 'in the past month...' followed by a question relating to experiences of stress, e.g., 'How often have you been upset by something that happened unexpectedly?' | Higher scores indicate more frequently experienced stress. |

*Note.* All surveys use computer adaptive testing so there is no set number of items.

### 2.3. Neuroimaging Measures

#### 2.3.1. MRI Acquisition and Pre-Processing

The acquisition parameters and processing steps of the HCP-YA are described in detail in Glasser et al., (2013). In brief, the HCP structural imaging protocol used a 3T Siemens Skyra scanner with a 32-channel head coil to acquire T1 and T2-weighted images. T1-weighted images were acquired with a 3D Magnetisation Prepared Rapid Acquisition Gradient Echo (3D MPRAGE) sequence, with 0.7 mm isotropic voxels, repetition time (TR) of 2400 ms, echo time (TE) of 2.14 ms, a field of view of 224 by 224 mm^2^, 256 sagittal slices, and an acquisition time of 7.40 minutes (Glasser et al., 2013). T2-weighted images were acquired with a 3D SPACE sequence with the same geometry as the T1-weighted images but a TR of 3200 ms, TE of 565 ms, and an acquisition time of 8.24 minutes. To calculate the cortical volume of each region, the HCP-YA team used FreeSurfer 5.3-HCP (Fischl, 2012; Glasser et al., 2013). The FreeSurfer pipeline, as detailed in Glasser et al. (2013), included co-registering and averaging T1 and T2-weighted images to improve pial surface quality, bias field correction, segmentation, and surface reconstruction.

#### 2.3.2. Regions of Interest

Cortical volume was extracted from 12 regions of interest (ROIs; depicted in Figure 1), parcellated based on the Desikan-Killiany atlas (Desikan et al., 2006). Cortical volume was selected as it has been found to be more sensitive to detecting inter-layer edges than cortical thickness or surface area (Simpson-Kent et al., 2021). The ROIs were selected based on a selection of regions in prior work by Metzler-Baddeley et al. (2016), which investigated working memory using regions from the central executive and salience networks. Additional studies (e.g., Cromheeke & Mueller, 2014; Diener et al., 2012; Kraljević et al., 2021; Menon, 2011) offer further support for the involvement of some of these regions in cognition and/or emotion. Supplementary Table 1 provides detailed justification for each region’s inclusion, including references to studies that have implicated the region in relevant psychological functions. Nine ROIs are from the central executive network and include the superior frontal gyrus, caudal middle frontal gyrus, rostral middle frontal gyrus, pars orbitalis, pars triangularis, pars opercularis, superior parietal gyrus, inferior parietal gyrus, and supramarginal gyrus. Three ROIs are from the salience network and include the insula, rostral anterior cingulate gyrus, and the caudal anterior cingulate gyrus. See Table 3 for the complete list of nodes included in the present study, including their abbreviations (used in figures) and network layers.

**Figure 1.**
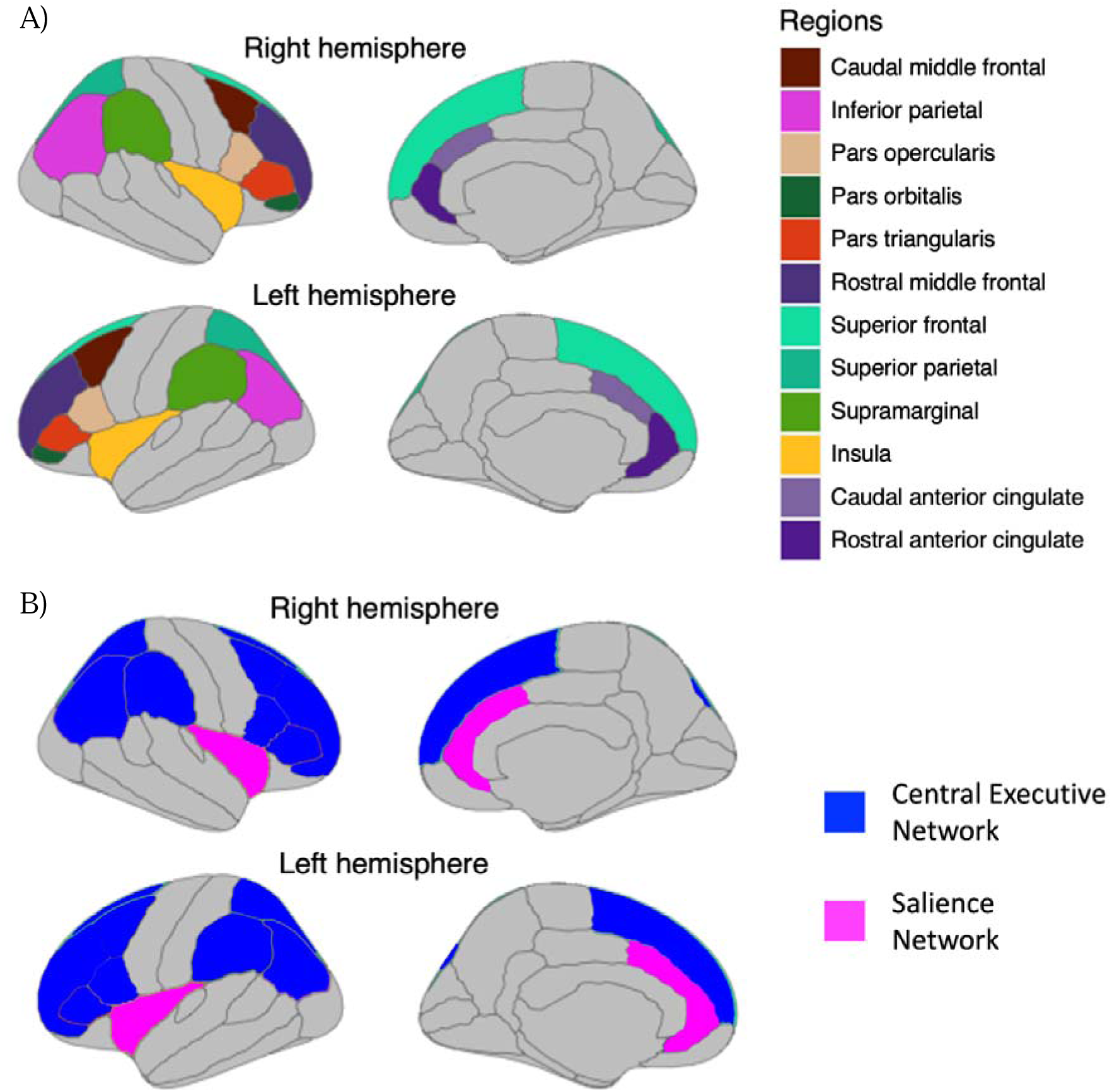
(A) Grey matter ROIs based on the Desikan-Killiany atlas; depicted bilaterally. (B) ROIs according to their network.

**Table 3.** Complete list of nodes in multilayer networks, including their abbreviation and layer.

| Node | Abbreviation | Layer |
| --- | --- | --- |
| Episodic Memory | Episodic | Cognition |
| Working Memory | Working | Cognition |
| Executive Function | Executive | Cognition |
| Attention | Attention | Cognition |
| Reading | Reading | Cognition |
| Vocabulary | Vocabulary | Cognition |
| Processing Speed | Speed | Cognition |
| Anger | Anger | Emotion |
| Fear | Fear | Emotion |
| Sadness | Sadness | Emotion |
| Stress | Stress | Emotion |
| Superior Frontal Gyrus | SF | Brain |
| Caudal Middle Frontal Gyrus | CMF | Brain |
| Rostral Middle Frontal Gyrus | RMF | Brain |
| Pars Orbitalis | POr | Brain |
| Pars Triangularis | PT | Brain |
| Pars Opercularis | POp | Brain |
| Superior Parietal Gyrus | SP | Brain |
| Inferior Parietal Gyrus | IP | Brain |
| Supramarginal Gyrus | SM | Brain |
| Insula | I | Brain |
| Rostral Anterior Cingulate | RAC | Brain |
| Caudal Anterior Cingulate | CAC | Brain |

### 2.4. Statistical Analysis

#### 2.4.1. Network Estimation

Figure 2 depicts the analysis pipeline used to conduct the multilayer network analyses. All network analyses and visualisations were completed using the *bootnet* wraparound package (Epskamp et al., 2018; version 1.5) in R (R Core Team, 2020; version 4.1.2) and the code to run these analyses can be found on the Open Science Framework website (https://osf.io/fj68r/). A psychometric bi-layer network was modelled for cognition and emotion variables, including seven cognitive nodes and four emotion nodes. This network used (Pearson) partial correlation networks with model selection via graphical lasso regularisation (Friedman et al., 2008) and tuning of the extended Bayesian information criterion (EBIC; J. Chen & Chen, 2008). Partial correlation networks include undirected edges between nodes that are weighted by the strength of the partial correlation between the two nodes, after controlling for all other nodes in the network.

**Figure 2.**
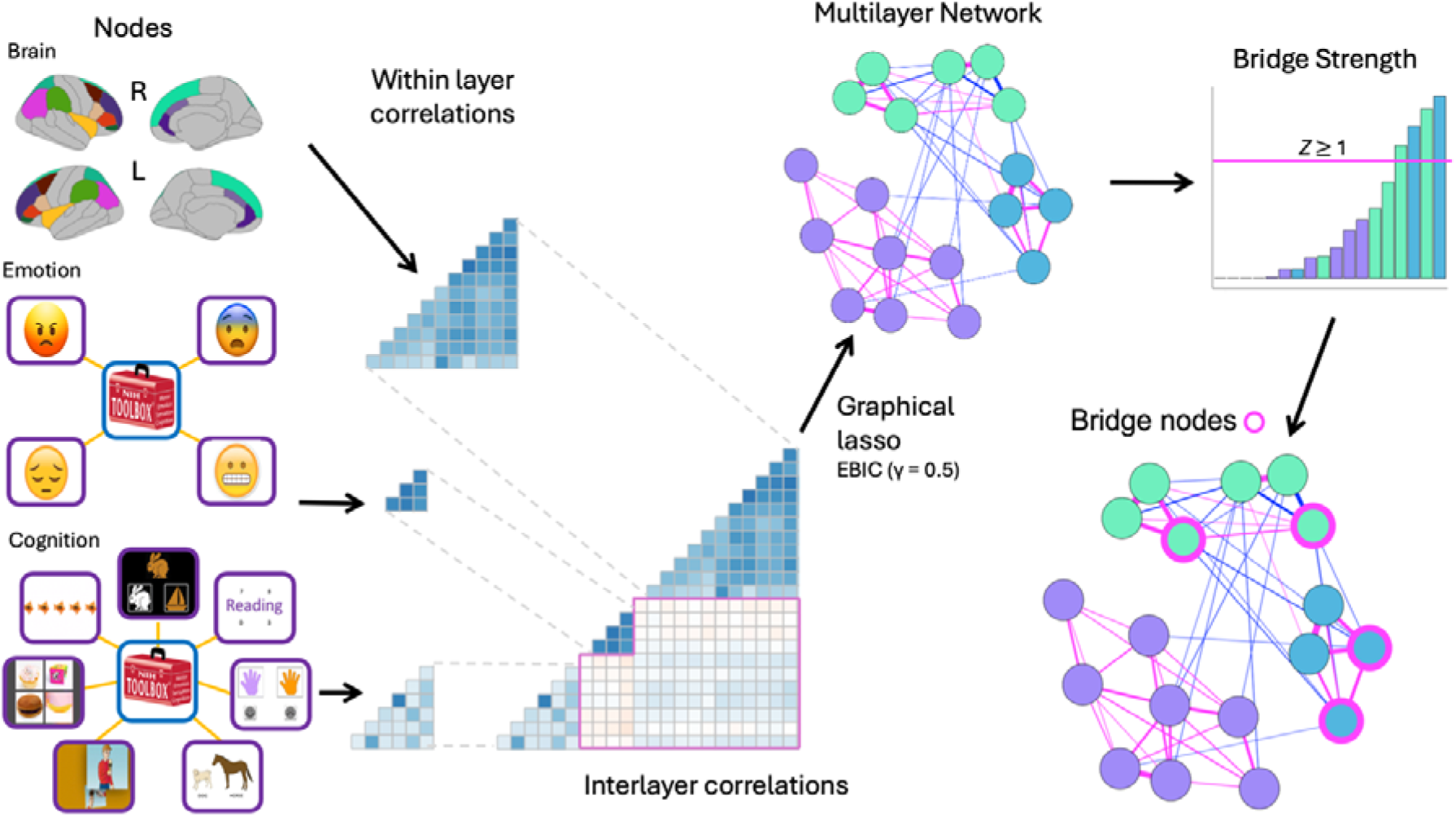
Overview of the analysis steps to generate multilayer networks and identify bridge nodes. First, individual variables for each layer (i.e., cognition, emotion and brain) were entered into a single correlation matrix. Graphical lasso model selection was then applied to generate the multilayer network from this matrix. Bridge strength was subsequently used to determine nodes that were important bridges between layers. “R” = right hemisphere, “L” = left hemisphere. “*Z*” *= z-*score. Note: This figure pertains specifically to the neuro-psychometric tri-layer network, while the psychometric bi-layer network followed the same steps but did not include the brain data.

Nonparanormal transformations were performed prior to calculating the network models using the *npn* option within the *estimateNetwork* function of *bootnet*. Transformation was necessary to address the non-normal distribution of most variables, ensuring appropriate model selection. Graphical lasso regularisation was used because it is the most conservative regularisation method for reducing the inclusion of spurious correlations in the network model. The EBIC hyperparameter was set to 0.5, to further reduce the likelihood of spurious correlations (Epskamp & Fried, 2018). Networks were visualised using the *qgraph* package (Epskamp et al., 2012; version 1.9.2), with node positioning determined by the Fruchterman-Reingold algorithm, which arranges strongly connected nodes near each other (Fruchterman & Reingold, 1991).

A neuro-psychometric tri-layer network was then modelled to include the 12 ROIs in addition to the cognitive and emotion variables. Each ROI included two nodes in the network, one for the cortical volume of each hemisphere i.e., 24 brain nodes in total. The tri-layer network was calculated in the same manner as the bi-layer network.

#### 2.4.2. Network Description

A matrix of zero-order Pearson correlations was plotted for the tri-layer network and is included in the supplementary materials (a section of this matrix also represents the zero-order correlations for the bi-layer network). These values were computed following nonparanormal transformation to align with the input data used in network estimation. The matrix is included to show the correlation values before conversion to partial correlations and regularisation, as well as to facilitate comparison to previous multilayer network studies (e.g., Simpson-Kent, 2021). The edge weights in the estimated networks reported in the Results section are the regularised partial correlations. The strongest edge weights are reported primarily for descriptive purposes, to illustrate the connectivity pattern within and between layers.

To describe the density of the networks, we calculated the percentage of edges present in the network from the total possible number of edges. Based on previous network descriptions, a network with less than 30% of possible edges would be considered low density, 30% to 50% of possible edges would be considered medium density, and more than 50% would be considered high density (Carmichael et al., 2023; Johal & Rhemtulla, 2023).

Bridge strength centrality was calculated to identify nodes that serve as bridges between layers (Jones et al., 2021). The *bridge* function of the *bootnet* package was used to calculate bridge centrality, after assigning nodes to pre-defined communities based on their layer, i.e., emotion, cognition, or brain layer. Bridge strength centrality was calculated for each node by summing the absolute edge weights between that node and nodes in different layers. In addition, normalisation was used to control for differences in community size, by dividing each node’s bridge strength value by the node’s maximum possible bridge strength value. Finally, bridge strength was calculated as a centrality coefficient and then standardised into a *z*-score. In line with previous work, bridge nodes were defined as those with a positive bridge strength centrality *z*-score equal to or greater than one (Simpson-Kent et al., 2021).

#### 2.4.3. Inspection of Edge Weights

For both networks, we inspected edge weights to characterise the pattern of within- and between-layer connections. This process included assessing the relative strength of edges, their predominant direction (positive or negative), and identifying the strongest edges in each layer. In addition, we examined the edge weights connected to bridge nodes to determine the number and strength of the bridge node’s inter-layer connections, aiding interpretation of how these nodes function as bridges between layers.

#### 2.4.4. Edge Weight and Bridge Strength Stability

To determine the reliability of edge weight estimates in both the bi-layer and tri-layer networks, non-parametric bootstrapping was implemented using *bootnet*, as described in Epskamp et al. (2018), with 2000 bootstraps. Edges that occurred in fewer than 50% of the bootstrapped samples require caution when being interpreted (Carmichael et al., 2023). For bridge strength estimates, correlation stability (CS) coefficients were calculated using case dropping bootstrapping (Epskamp et al., 2018). These coefficients reflect the maximum proportion of cases that can be removed while maintaining a correlation of at least 0.7 with the original centrality estimates, with 95% probability. Simulation studies conducted by Epskamp et al. (2018) suggest that CS coefficients below 0.25 indicate that the rank order of centrality indices is unstable and should not be interpreted. CS coefficients above 0.25 are considered stable, however, if the CS coefficient is below 0.5 then more caution is needed in interpretation.

## 3. Results

3.1. **Psychometric Bi-layer Network: Description and Edge Weight Stability**

The bi-layer network (Figure 3) had high density as it included 31 edges out of a possible 55 (56%), with a mean edge weight of 0.127 (median = 0.065). The majority (27/31) of the edges were positive and ranged from 0.002 (between Fear and Reading) to 0.601 (between Reading and Vocabulary). The strongest edges were between nodes within the same layer, i.e., between Reading and Vocabulary (0.601), Attention and Executive (0.391), and Fear and Sadness (0.366).

**Figure 3.**
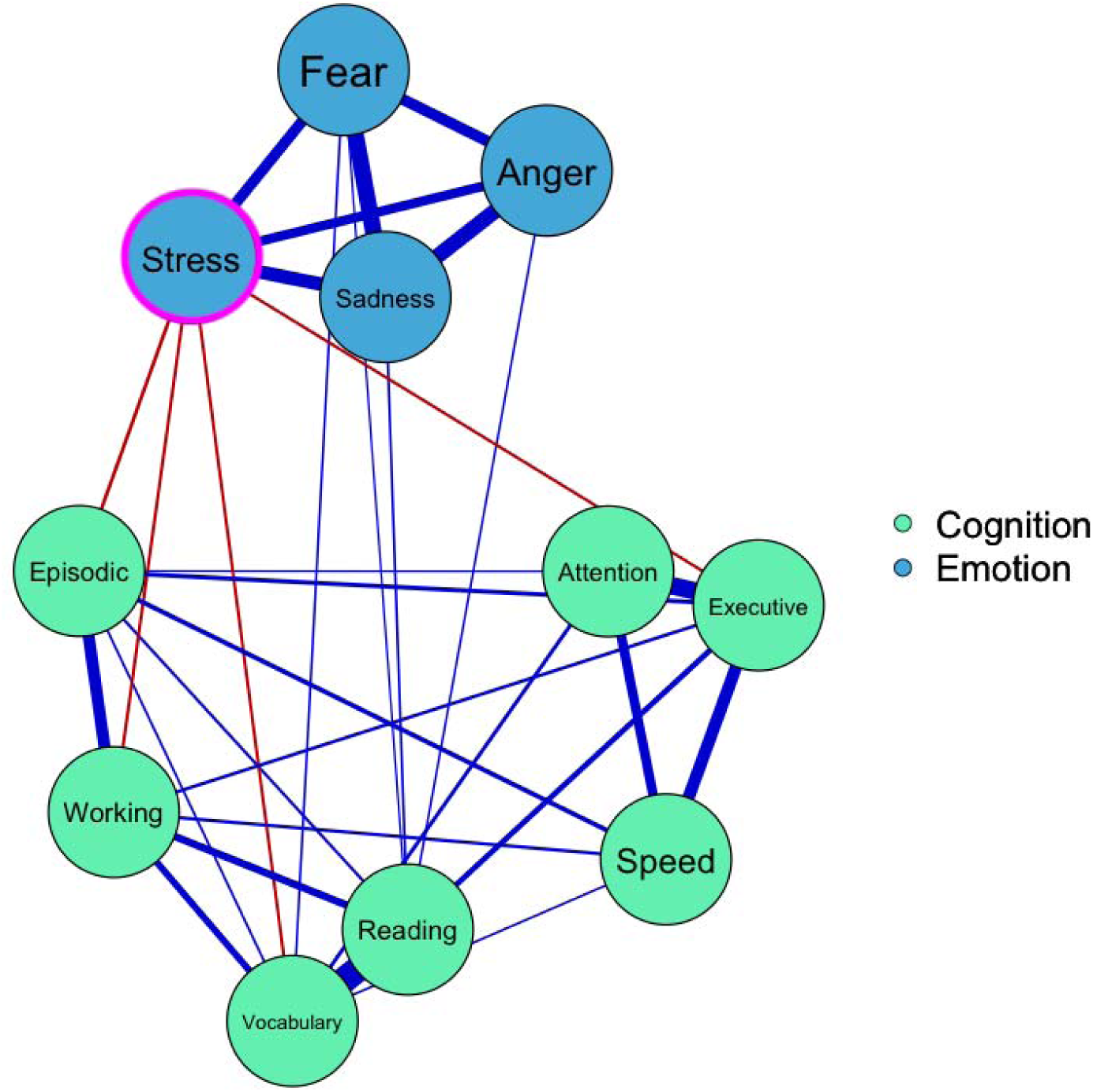
Bi-layer network (*n* = 1109). Nodes (circles) are variables from the NIH Toolbox Cognition Battery (cognition; colour coded green) and NIH Toolbox Emotion Battery (emotion; colour coded blue). Undirected edges (lines between nodes) represent partial correlations between variables, after controlling for all other variables, with thicker lines representing stronger relationships. Blue edges represent positive relationships, whereas red edges represent negative relationships. A magenta circle indicates the node with a bridge centrality *z*-score ≥ 1. Abbreviations for node labels are defined in Table 3.

There were eight inter-layer edges. Four were positive: Reading and Sadness (0.016), Fear and Vocabulary (0.012), Anger and Reading (0.009), and Fear and Reading (0.002). The remaining four were negative, all involving Stress: Episodic (-0.043), Working (-0.032), Executive (-0.030) and Vocabulary (-0.029)

Four edges should be interpreted with caution, as they were found in less than 50% of the 2000 bootstrapped samples: Fear and Vocabulary (40%), Fear and Reading (42%), Anger and Reading (44%), and Attention and Episodic (45%). All of the other edges in the bi-layer network can be considered stable, as they were found in more than 50% of the bootstrapped samples.

### 3.2. Psychometric Bi-layer Network: Bridge Strength Centrality

One node, stress (*z* = 2.211), was identified as having a *z*-score greater than one standard deviation (Figure 4) and was therefore the only bridge node in the bi-layer network. As noted above, Stress had inter-layer edges with Episodic, Working, Executive and Vocabulary. Supplementary Table 2 shows raw, normalised, and *z*-scored (after normalisation) centrality values for all nodes. Regarding stability, bridge strength had a CS coefficient of 0.361.

**Figure 4.**
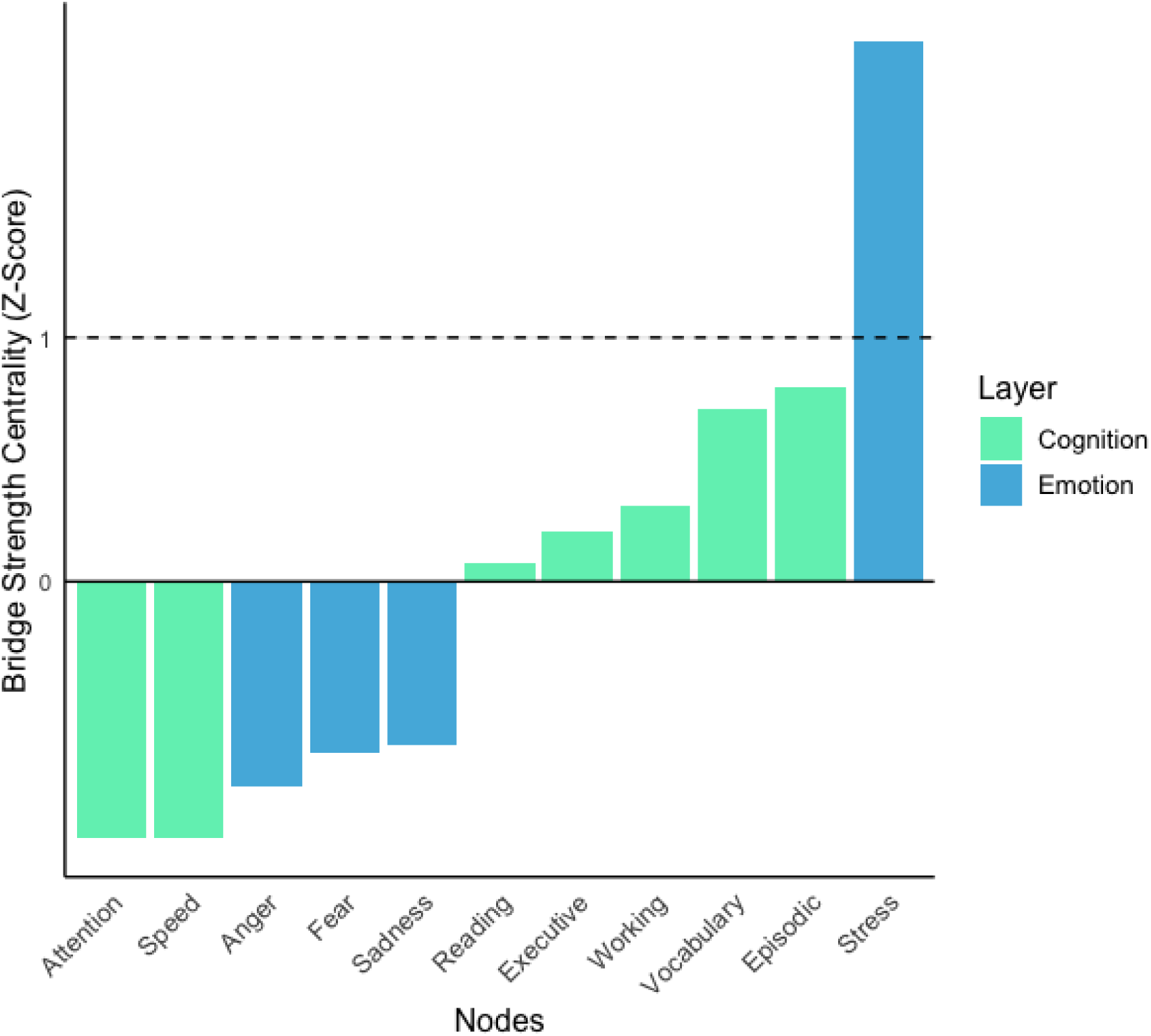
Bar graph of bridge strength centrality scores (*n* =1109). The dashed line indicates a *z*-score of 1, nodes at or above this line are interpreted as bridge nodes. Abbreviations for node labels are defined in Table 3.

**3.3. Neuro-Psychometric Tri-layer Network: Description and Edge Weight Stability** The tri-layer network (Figure 5; see Supplementary Figure 1 for the zero-order correlation matrix) had medium density as it included 243 edges out of a possible 595 (41%), with a mean edge weight of 0.062 (median = 0.030). As with the bi-layer network, the majority of edge weights were positive (214/ 243), and the strongest edges were within layers. The strongest edge was within the brain layer between the left and right I (0.594), followed by the strongest edge within the cognition layer between Reading and Vocabulary (0.591). This was followed by the edges between the left and right hemispheres of each of the following regions: SP (0.517), RMF (0.496), IP (0.472), and SF (0.445). The strongest edge between emotion nodes was between Fear and Sadness (0.366). The strongest negative edges were also within layers, the strongest of which was between left CAC and left SF (-0.232).

**Figure 5.**
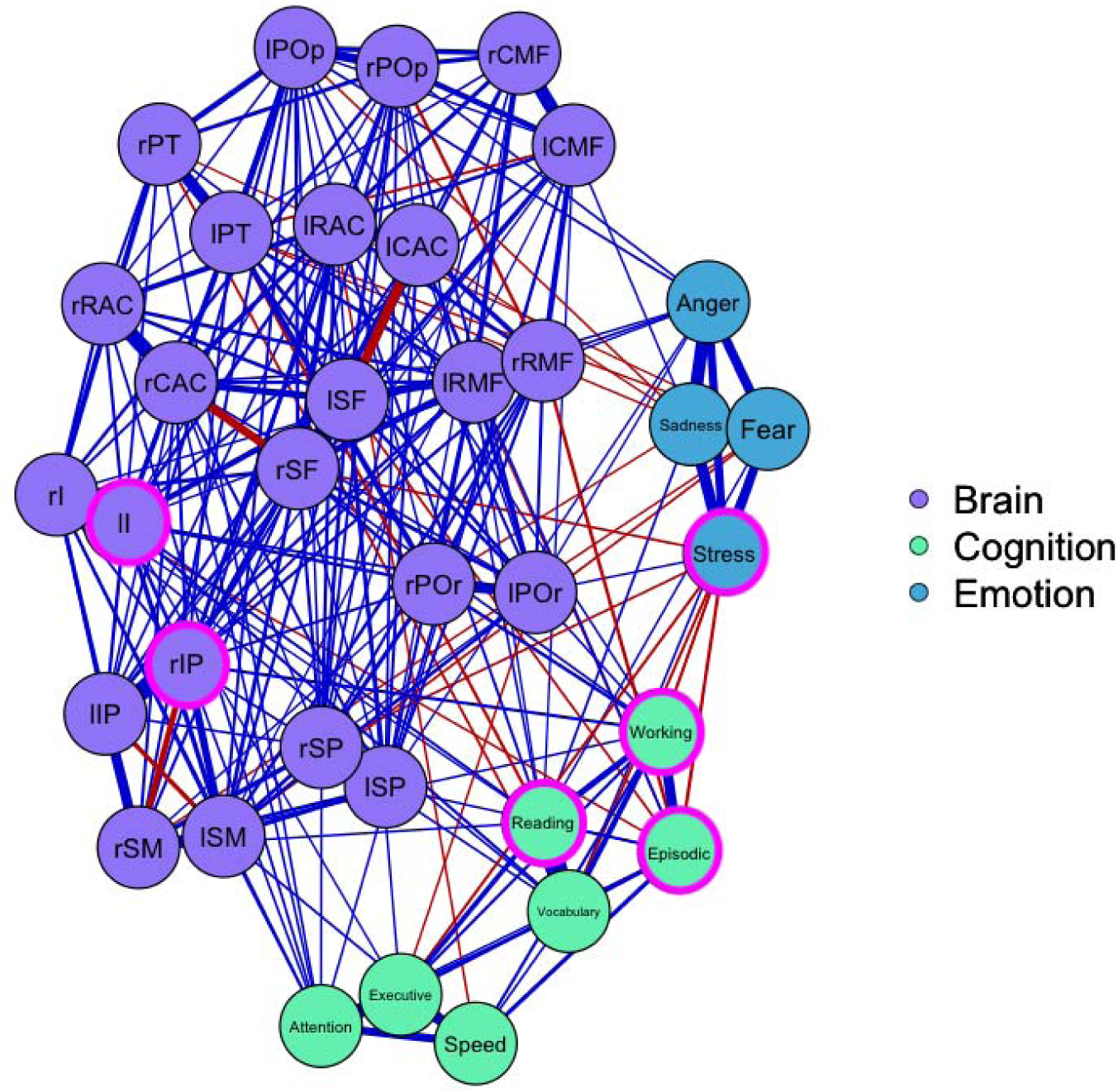
Tri-layer network (*n* = 1109). Nodes (circles) are variables from the NIH Toolbox Cognition Battery (cognition; colour coded green), NIH Toolbox Emotion Battery (emotion; colour coded blue), and the cortical volume of ROIs from the central executive and salience networks (Brain; colour coded purple). Undirected edges (lines between nodes) represent partial correlations between variables, after controlling for all other variables, with thicker lines representing stronger relationships. Blue edges represent positive relationships, whereas red edges represent negative relationships. Magenta circles around nodes indicate that the node has a bridge centrality *z*-score ≥ 1. “l” before a ROI indicates the left hemisphere, “r” before a ROI indicates the right hemisphere. Abbreviations for node labels are defined in Table 3.

There were 63 inter-layer edges. Forty-one of these edges were positive and the strongest was between Working and right POr (0.047), followed by Working and right IP (0.041), and Vocabulary and left I (0.033). There were 22 negative inter-layer edges, the strongest of which was between Episodic and right RMF (-0.046), followed by Episodic and Stress (-0.045), Reading and left PT (-0.036), Working and Stress (-0.029), and Executive and Stress (-0.029).

The majority of edge weights in the tri-layer network were stable when comparing edge weights across 2000 bootstrapped samples. However, 43 (17.70%) of the 243 edges were included in less than 50% of the bootstrapped samples and should be interpreted with caution (these edges are listed in Supplementary Table 3).

### 3.4. Neuro-Psychometric Tri-layer Network: Bridge Strength Centrality

For the tri-layer network, there were six bridge nodes, having *z*-scores greater than one standard deviation (Figure 6). Two of these nodes were brain nodes, three were cognitive nodes and one was an emotion node. In descending order, the bridge nodes were left I (*z* = 1.845), Working (*z* = 1.467), Stress (*z* =1.371), Episodic (*z* = 1.252), Reading (*z* = 1.226), and right IP (*z* =1.073). Supplementary Table 4 shows raw, normalised, and *z*-scored (after normalisation) centrality values for all nodes. Regarding stability, bridge strength had a CS coefficient of 0.438.

**Figure 6.**
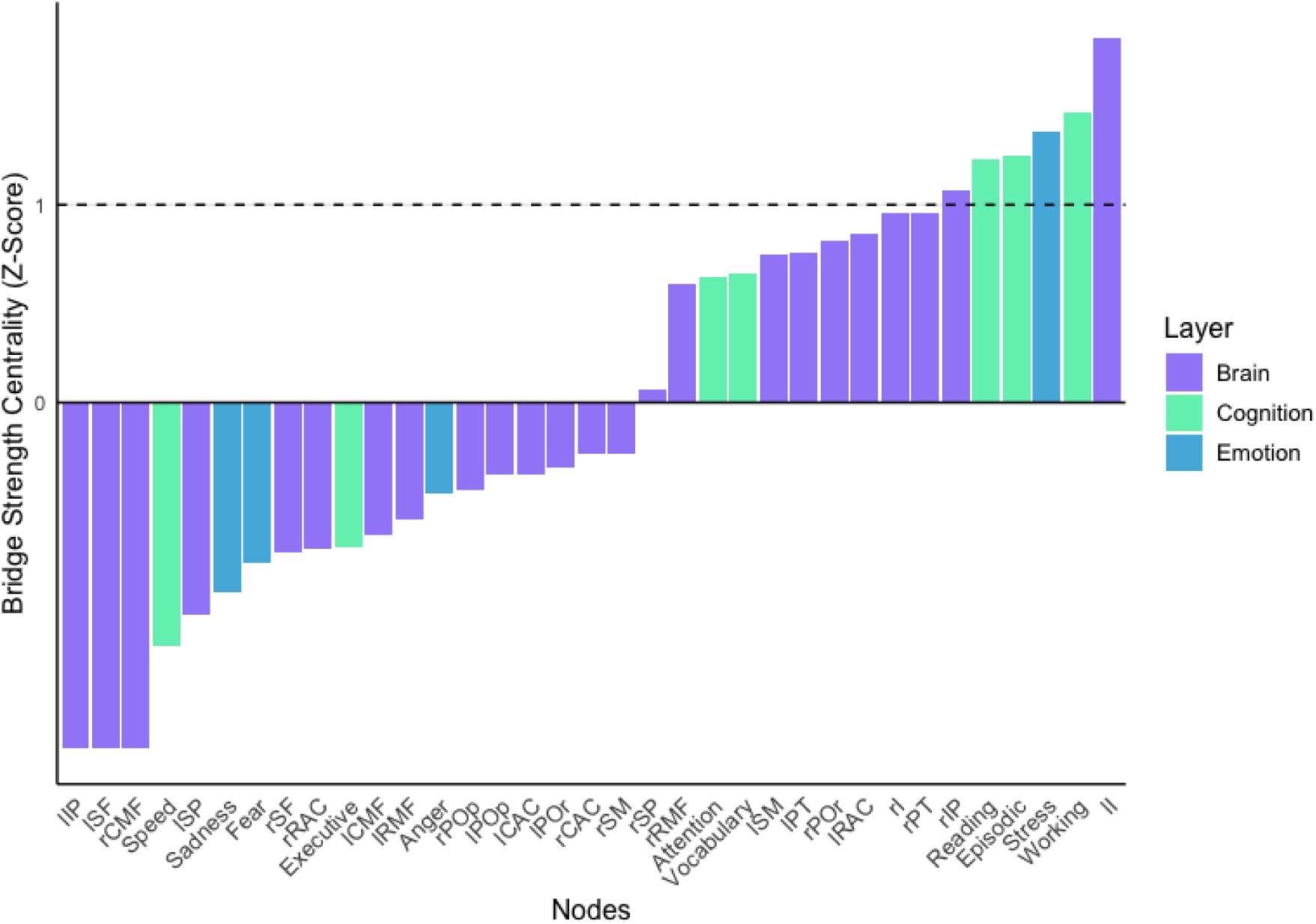
Bar graph of bridge strength centrality scores (*n* =1109). The dashed line indicates a *z*-score of 1, nodes at or above this line are interpreted as bridge nodes. “l” before a ROI indicates the left hemisphere, “r” before a ROI indicates the right hemisphere. Abbreviations for node labels are defined in Table 3.

The left I had inter-layer edges with four nodes, i.e., Vocabulary (0.033), Reading (0.025), Anger (0.007), and Executive (0.005). Working had inter-layer edges with eight nodes, i.e., right POr (0.047), right IP (0.041), right CAC (0.014), right SM (0.011), left POr (0.010), left RAC (0.008), Fear (-0.001), and Stress (-0.029). Stress had inter-layer edges with seven nodes, i.e., left POr (0.008), right SP (-0.015), right SF (-0.019), Vocabulary (-0.028), Executive (-0.029), Working (-0.029), and Episodic (-0.045). Episodic had inter-layer edges with six nodes, i.e., left CAC (0.025), left SP (0.009), right I (-0.009), right PT (-0.016), Stress (-0.045), and right RMF (-0.046). Reading had inter-layer edges with ten nodes, i.e., left I (0.025), left RAC (0.024), Sadness (0.018), right POp (0.014), right SM (0.012), Fear (0.007), left RMF (0.005), left CMF (0.005), right I (0.004), and left PT (-0.004). Right IP had inter-layer edges with two variables, i.e., Working (0.041) and Attention (0.014).

## 4. Discussion

The present study aimed to explore the complex relationships between cognition, emotion, and brain structure in healthy young adults, utilising a novel analytic framework. Specifically, the aim was to use a multilayer network approach to identify nodes that are central to the relationship between cognition and emotion, both in a purely psychometric context, as well as when integrating neuroscientific data. In the psychometric bi-layer network of cognition and emotion, it was found that stress is the only bridge node. Furthermore, in the neuro-psychometric tri-layer network consisting of cognition, emotion and brain regions, it was found that the left insula was the most central bridge node overall, working memory was the most central cognition bridge node, and stress was the most central emotion bridge node. In addition, episodic memory, reading and the right inferior parietal emerged as bridge nodes. The following discussion will focus on each of these bridge nodes.

In the bi-layer network, the only bridge node was stress, which was negatively related to episodic memory, working memory, executive function, and vocabulary. This finding aligns with an extensive body of research on stress and cognition (Calvo & Gutiérrez-García, 2016; Sandi, 2013). Consistent with the present study’s finding that the strongest edge across layers is between stress and episodic memory, Zaheed et al. (2021) found that perceived stress was associated with worse episodic memory in healthy older adults (aged 60 to 95 years). Stress has also been shown to impair both working memory and executive function (Shields et al., 2016). A meta-analysis by Shields et al. proposes that stress impairs executive function and working memory via the interplay of multiple hormones and immune system processes, as well as psychosocial factors. The effect of stress on memory may in turn explain our finding of a negative relationship between stress and vocabulary. It has been suggested that students who experience more stress during test taking are more likely to have difficulty retrieving semantic memories, even when the relevant knowledge is intact (Vogel & Schwabe, 2016). Alternatively, early-life stress may also play a role. For example, children’s vocabulary development has been shown to be negatively affected by parental stress (Deniz Can & Ginsburg-Block, 2021), suggesting that individuals exposed to high-stress environments during development may carry forward both increased susceptibility to stress and reduced vocabulary performance into adulthood.

Stress was also identified as the most central bridge node in the emotion layer of the tri-layer network, where it was connected to the same cognitive variables as in the bi-layer network, and additionally showed negative associations with the right superior parietal and right superior frontal gyri, and a positive association with the left pars orbitalis. The negative associations observed with the right superior parietal and right superior frontal gyri align with findings from a large-scale neuroimaging mega-analysis of individuals with post-traumatic stress disorder, which reported reduced cortical volume in these right-lateralised regions (Wang et al., 2021). Our findings are consistent with the notion that chronic stress contributes to volume reductions in parietal and frontal regions, particularly in the right hemisphere. The positive association between stress and volume in the left pars orbitalis is more difficult to interpret, as relatively few studies have examined structural correlates of stress in this region. One relevant study by Lehne et al. (2014) found that the left pars orbitalis was activated during the experience of musical tension, suggesting a role in the subjective appraisal or awareness of emotional tension. It is possible that this region supports metacognitive awareness of one’s own stress levels or internal states. In this context, individuals with greater volume in the left pars orbitalis may also exhibit greater subjective sensitivity to stress, which could lead them to report higher levels of perceived stress. While this interpretation remains speculative, it may offer a meaningful link between structural differences in this region and the self-reported stress levels observed in the present study. Together, the results of the bi-layer and tri-layer networks underscore the centrality of stress in linking emotion and cognition, and demonstrate the utility of neuro-psychometric network models for identifying potential brain substrates of this relationship.

In the tri-layer network, the left insula was identified as the most central bridge node, connecting both emotion and cognition. Specifically, it showed connections with anger as well as vocabulary, reading, and executive function. This finding aligns with prior research highlighting the insula’s integral role in integrating emotional and cognitive processes (Gasquoine, 2014). It has been reported that the insula is crucial to interoception, which in turn gives rise to emotional feeling (Damasio, 2003). The insula’s role in cognition may be more indirect; while insular lesions can be associated with cognitive impairment, these effects are often attributed to diaschisis or disconnection rather than direct involvement in cognitive processing per se (Gasquoine, 2014). Supporting the insula’s direct involvement in emotion and downstream involvement in cognition is a brain network model proposed by Menon and Uddin (2010), who suggest that the insula is activated in relation to salient events and then signals to other brain regions to facilitate working memory and attention, while generating the emotional feelings appropriate to that event. Given its role in both cognitive and emotional integration, the insula may represent a promising target for interventions aimed at enhancing these domains, for example through mindfulness meditation (Sharp et al., 2018).

Although the insula is frequently implicated in stress-related processing, particularly through its role in the salience network during acute stress (Berretz et al., 2021), our network did not reveal a direct connection between stress and the insula. This may reflect an indirect relationship mediated by intermediary cognitive and emotional nodes. Alternatively, our findings may reflect that the insula is less involved in chronic stress. Much research on the role of the insula in stress focuses on acute physiological or experimentally induced stress responses (Berretz et al., 2021), whereas the self-reported measure for stress included in the present study asked participants to reflect on their stress over the previous month.

Importantly, the insula is not a uniform structure, and recent parcellation work has demonstrated clear functional subdivisions (Chang et al., 2013). Chang et al., using resting-state connectivity and meta-analytic coactivation, identified a tripartite division: a ventroanterior subdivision associated with affective and autonomic processing; a dorsoanterior subdivision associated with higher cognitive control and executive function; and a posterior subdivision related to sensorimotor processing. While the present analysis, based on cortical volume from the Desikan-Killiany atlas, did not differentiate between these subregions, future multilayer network studies may benefit from incorporating finer-grained parcellations of the insula to examine whether specific subdivisions differentially contribute to bridging cognition and emotion.

Working memory was the most central bridge node in the cognition layer of the tri-layer network. Just like in the bi-layer network, working memory was negatively associated with stress. However, the increased importance of working memory in the tri-layer network compared to the bi-layer network is due to its many edges with brain region nodes. Working memory had edges with six brain region nodes, specifically the left and right pars orbitalis, left rostral anterior cingulate, right caudal anterior cingulate, right inferior parietal gyrus, and right supramarginal gyrus. Many of these regions have previously been shown to exhibit structural changes following working memory training (Metzler-Baddeley et al., 2016). Given that working memory was central only in the tri-layer network, and not in the bi-layer network, these findings support the added value of incorporating neuroscientific data when aiming to identify key psychometric variables within a network, potentially allowing researchers to better target future interventions (Bathelt et al., 2022; Blanken et al., 2021; Hilland et al., 2020; Simpson-Kent et al., 2021). These findings also highlight the potential utility of working memory training programs, such as that implemented in Metzler-Baddeley et al. (2016), which may not only improve working memory but also have downstream benefits for broader cognitive functioning, while potentially reducing negative emotional states.

Episodic memory was the second most central bridge node in the cognition layer of the tri-layer network. Episodic memory was also found to be a central node in Ferguson’s (2021) psychometric network in healthy older adults. While Ferguson’s psychometric network only included cognitive abilities, our findings show that episodic memory is not only important in the network of cognitive abilities, but it also bridges cognition with emotion and brain structure. Furthermore, the participants in Ferguson’s study had a mean age of 76 years, therefore, the present study’s results in a young adult sample suggest that episodic memory’s centrality may be stable across the lifespan in those without a neurodegenerative condition. As Ferguson compared the healthy network to an Alzheimer’s Disease network and found that processing speed became more central in the Alzheimer’s Disease network, future research could compare the present findings to a network of cognition, emotion and brain structure in neurodegenerative conditions. Such comparisons may reveal different patterns of edges and bridge nodes than the present study’s network of healthy young adults.

The third most central bridge node in the cognition layer of the tri-layer network was reading. This aligns with findings from Simpson-Kent et al. (2021), who identified reading as the most central bridge node in their tri-layer network of cognition, cortical volume, and fractional anisotropy. The prominence of reading in both studies supports the view that reading ability relies on the integration of multiple brain systems (Price, 2012). Notably, our results also suggest that reading is not only cognitively central, but also functionally linked to emotion, as reflected in its positive associations with sadness and fear. These connections may reflect overlapping mechanisms between emotional sensitivity and verbal or linguistic processing, though this relationship warrants further investigation.

The right inferior parietal gyrus was the second most central bridge node in the brain layer, with interlayer edges to working memory and attention. This finding aligns with previous research examining the inferior parietal gyrus’ involvement in cognitive tasks (Cao et al., 2021; Ren et al., 2019). For example, Cao et al. found that the inferior parietal gyrus was activated during a 2-back task in both healthy controls and individuals with major depressive disorder. They suggested that the inferior parietal gyrus is involved in focusing attention on external stimuli, which in turn facilitates working memory. Similarly, a relationship between the right inferior parietal gyrus and attention has been found in virtual lesion studies using Transcranial Magnetic Stimulation (TMS; Bareham et al., 2018; Karhson et al., 2015). Our findings suggest that the right inferior parietal gyrus may represent a promising target for intervention. For example, stimulation of this region via therapeutic TMS may enhance working memory and attention. Improving cognitive function in this way could also indirectly support better stress regulation, given the observed interplay between these cognitive and emotional processes in the present network.

A notable absence from the bridge nodes in the tri-layer network was the superior frontal gyrus. The left superior frontal gyrus was found to be associated with both cognition and emotion in the study of Kraljević et al., (2021) using HCP-YA data. Furthermore, Simpson-Kent at al. (2021) found the bilateral superior frontal gyrus to be central to their network of cognition and cortical volume. However, Simpson-Kent et al. used community-based bridge centrality, in which there was more than one community that included brain regions. This approach may have elevated the importance of brain regions that bridge to other brain regions rather than those specifically connected to cognition. Furthermore, although Kraljević et al., (2021) extracted similar cognition, emotion and brain imaging data as the present study, they used multiple linear regression models, suggesting that the present study’s use of partial correlation networks may have revealed that the superior frontal gyrus’ relationship to cognition and emotion is mediated by its relationship to other brain regions, indicating a more indirect role.

The findings of the present study should be interpreted cautiously given the exploratory and novel nature of the analysis. Firstly, the rank order of bridge nodes in both networks demonstrated stability; however, based on the correlation stability (CS) thresholds proposed by Epskamp et al. (2018), this order may need to be interpreted with some caution. It is worth noting that the CS values reported here are comparable to those observed in a prior multilayer network study by Simpson-Kent et al. (2021). Importantly, Epskamp et al.’s guidelines were developed for single-layer networks and strength centrality, not for multilayer networks or bridge strength. It remains unclear how appropriate these thresholds are when applied to bridge strength estimates in multilayer networks, particularly those that integrate psychometric and neuroimaging data. In such networks, bridge strength reflects only inter-layer edges, which in general are likely to be weaker and less stable, given the nature of exploring statistical relationships across domains. Considering that the present study involved a relatively large sample, it is likely that detecting bridge nodes with sufficient stability in similar multilayer designs will require comparable or even larger sample sizes.

A further limitation of this study is the inclusion of only one neuroimaging modality: cortical grey matter volume. While this decision was based on evidence supporting structural covariance across cortical regions (Alexander-Bloch et al., 2013) and its demonstrated sensitivity to inter-layer edges in prior multilayer network studies (Simpson-Kent et al., 2021), it necessarily limits the scope of neural characteristics examined. Other grey matter features such as cortical thickness or surface area, and white matter measures such as fractional anisotropy or metrics from fixel-based analysis (Dhollander et al., 2021), may provide complementary insights. However, integrating additional modalities would require a more complex multilayer network with additional layers and nodes, which could impair interpretability. In particular, as nodes become connected across a larger number of layers, bridge strength may become diluted across many weak inter-layer connections, making it more difficult to identify specific cross-domain processes or intervention targets.

To conclude, this is the first study to integrate a psychometric network of both cognition and emotion with a structural covariance network of cortical brain regions. Specifically, using a large sample of healthy young adults, this multilayer network analysis revealed that the left insula is central to the relationship between the brain, cognition, and emotion. Furthermore, working memory and stress may be important psychological characteristics to consider for improving overall cognition and emotion in this population. Further advancement of this analytic approach may allow for comprehensive and targeted treatment strategies, as well as a greater understanding of the complex and dynamic relationship between the brain and the mind.

## Acknowledgments

The authors would like to thank the researchers of the Human Connectome Project, led by the WU-Minn Consortium (Principal Investigators: David Van Essen and Kamil Ugurbil; 1U54MH091657) as well as the participants who volunteered as part of the Young Adult data collection. In addition, we acknowledge the funding of this project by the 16 NIH Institutes and Centers that support the NIH Blueprint for Neuroscience Research; and by the McDonnell Center for Systems Neuroscience at Washington University. LFS is supported by a Deakin University Postgraduate Research scholarship.

