## Supplementary materials v2 for "The Left Insula Bridges Cognition, Emotion, and Brain Structure: A Multilayer Network Analysis of the Human Connectome Project-Young Adult"

**Supplementary Table 1**

Regions included in the tri-layer network with associated psychological functions

| Network | Cortical region | Associated psychological functions |
| --- | --- | --- |
| Central executive | Left superior frontal | Affect and cognition (Kraljević et al., 2021). Working memory (Boisgueheneuc et al., 2006) |
|  | Right superior frontal | Response inhibition (Hu et al., 2016) |
|  | Left caudal middle frontal | Detection of unfavourable outcomes (Ridderinkhof et al., 2004), cognitive control and emotional modulation (Diener et al., 2012) |
|  | Right caudal middle frontal | Working memory (Metzler-Baddeley et al., 2016), detection of unfavourable outcomes (Ridderinkhof et al., 2004), cognitive control and emotional modulation (Diener et al., 2012) |
|  | Left and right rostral middle frontal | Emotional regulation (Waugh et al., 2014) |
|  | Left and right pars orbitalis | Language processing (De Carli et al., 2007), inhibition (Wildgruber et al., 2005) |
|  | Left and right pars triangularis | Cognitive control, response inhibition, emotion-cognition interaction (Cromheeke & Mueller, 2014) |
|  | Left and right pars opercularis | Cognitive control, response inhibition, emotion-cognition interaction (Cromheeke & Mueller, 2014) |
|  | Left and right superior parietal | Manipulation of information in working memory (Koenigs et al., 2009). Attentional response switching (Rushworth et al., 2001) |
|  | Left and right inferior parietal | Attention to emotional/ social cues (Radua et al., 2010)  attention, language and social cognition (Numssen et al., 2021) |
|  | Left supramarginal | Phonological decisions (Hartwigsen et al., 2010) |
|  | Right supramarginal | Empathy (Silani et al., 2013). Phonological decisions (Hartwigsen et al., 2010) |
| Salience | Left insula | Emotional processing, empathy, anger, fear, sadness (Diener et al., 2012; Phan et al., 2002) |
|  | Right insula | Emotional processing, empathy, anger, fear, sadness (Diener et al., 2012; Phan et al., 2002). Working memory (Metzler-Baddeley et al., 2016) |
|  | Left and right rostral anterior cingulate | Emotional tasks with cognitive demand (Diener et al., 2012; Phan et al., 2002), internal emotional response and response selection (Devinsky et al., 1995) |
|  | Left and right caudal anterior cingulate | Emotional task with cognitive demand (Diener et al., 2012; Phan et al., 2002). Internal emotional response and response selection (Devinsky et al., 1995) |

**Supplementary Table 2**

Raw, normalised and standardised (*z*-scores after normalisation) bridge strength centrality values for the psychometric network.

| Node | Raw | Normalised | *Z*-Score |
| --- | --- | --- | --- |
| Stress | 0.13434856 | 0.01919265 | 2.21052679 |
| Episodic | 0.04337991 | 0.01084498 | 0.79351935 |
| Vocabulary | 0.04131229 | 0.01032807 | 0.70577523 |
| Working | 0.03205963 | 0.00801491 | 0.31311869 |
| Executive | 0.02951809 | 0.00737952 | 0.20526287 |
| Read | 0.0264987 | 0.00662467 | 0.07712806 |
| Sadness | 0.01573274 | 0.00224753 | -0.6658859 |
| Fear | 0.01413909 | 0.00201987 | -0.7045317 |
| Anger | 0.00854822 | 0.00122117 | -0.8401093 |
| Attention | 0 | 0 | -1.0474021 |
| Speed | 0 | 0 | -1.0474021 |

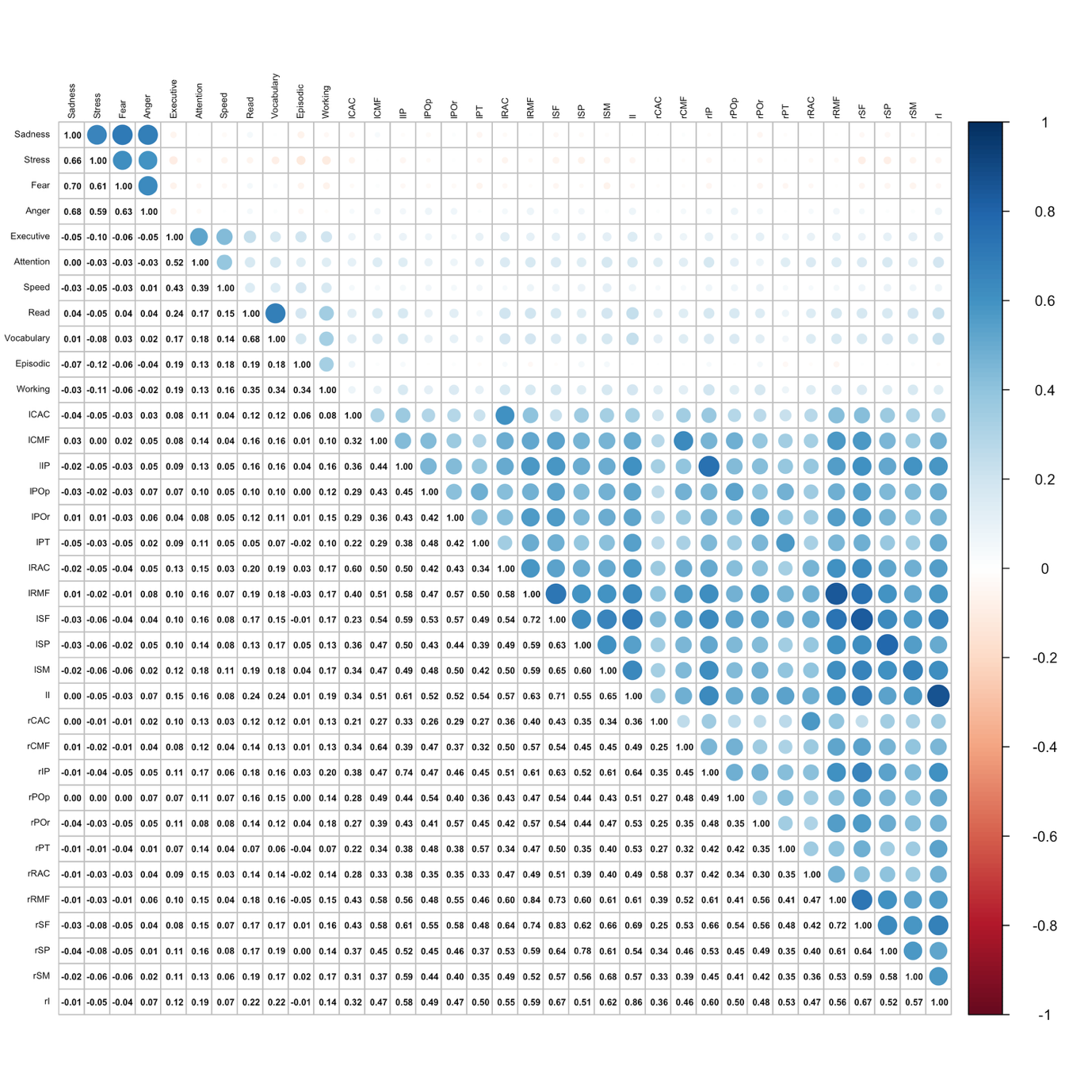

**Supplementary Figure 1.** Correlation matrix for cognition and emotion variables, and cortical volume of ROIs. Values represent nonparanormal transformed Pearson correlations. Blue circles indicate positive correlations, and red circles represent negative correlations. Circle size corresponds to the strength of the correlation. Lines separate inter-layer correlations from within-layer correlations. Values below and to the left of the green lines are the bi-layer inter-layer correlations (these are equivalent to those in the bi-layer correlation matrix, therefore the bi-layer correlation matrix is not included in the supplementary materials). Values below and to the left of the purple line are relationships between psychometric variables and brain structure.

**Supplementary Table 3**

Neuro-psychometric network edges found in less that 50% of bootstrapped samples

| Edge | % of bootstrapped samples |
| --- | --- |
| Left pars orbitalis - Right superior parietal | 49.45% |
| Working - Right caudal anterior cingulate | 49.40% |
| Read - Right pars opercularis | 49.30% |
| Left caudal anterior cingulate - Left rostral middle frontal | 49.15% |
| Left rostral anterior cingulate - Right pars opercularis | 48.80% |
| Attention - Left caudal middle frontal | 48.70% |
| Read - Right insula | 48.35% |
| Anger - Left insula | 48.20% |
| Right pars orbitalis - Right supramarginal | 47.60% |
| Left inferior parietal - Left superior frontal | 46.50% |
| Left caudal middle frontal - Left inferior parietal | 46.35% |
| Left caudal middle frontal - Left supramarginal | 46.15% |
| Fear - Vocabulary | 46.05% |
| Working - Right supramarginal | 46.00% |
| Right pars opercularis - Right rostral middle frontal | 44.25% |
| Episodic - Right insula | 43.80% |
| Executive - Left insula | 42.55% |
| Left caudal middle frontal - Left pars triangularis | 41.80% |
| Read - Left caudal middle frontal | 41.25% |
| Left pars triangularis - Right caudal anterior cingulate | 40.95% |
| Left caudal middle frontal - Right inferior parietal | 40.60% |
| Fear - Right supramarginal | 40.25% |
| Right inferior parietal - Right superior parietal | 39.85% |
| Sadness - Left caudal middle frontal | 39.55% |
| Anger - Left superior parietal | 38.95% |
| Executive - Left rostral anterior cingulate | 38.70% |
| Executive - Left pars orbitalis | 38.40% |
| Episodic - Left rostral middle frontal | 37.65% |
| Anger - Left pars orbitalis | 37.45% |
| Episodic - Left superior parietal | 37.20% |
| Left caudal anterior cingulate - Left pars orbitalis | 37.15% |
| Sadness - Left pars triangularis | 36.70% |
| Speed - Left rostral anterior cingulate | 35.40% |
| Fear - Left pars triangularis | 35.10% |
| Left superior parietal - Right inferior parietal | 34.90% |
| Read - Left rostral middle frontal | 32.35% |
| Sadness - Right pars orbitalis | 30.45% |
| Fear - Working | 30.25% |
| Stress - Left pars orbitalis | 28.45% |
| Sadness - Left caudal anterior cingulate | 27.70% |
| Sadness - Left pars opercularis | 27.45% |
| Left caudal anterior cingulate - Right pars opercularis | 27.20% |
| Fear - Right pars triangularis | 25.65% |

**Supplementary Table 4**

Raw, normalised and standardised (*z*-scores after normalisation) bridge strength centrality values for the neuro-psychometric network. Note: Raw scores do not take into account layer size and bias towards nodes in layers with fewer nodes, i.e., cognition and emotion layers. The order of the table is descending based on scores after normalisation to correspond with the bridge strength results in the main text.

| Node | Raw | Normalised | *Z*-Score |
| --- | --- | --- | --- |
| lI | 0.0705933 | 0.00641757 | 1.84482688 |
| Working | 0.16081893 | 0.00574353 | 1.46735266 |
| Stress | 0.17269417 | 0.00557078 | 1.37060758 |
| Episodic | 0.15003914 | 0.00535854 | 1.25174991 |
| Read | 0.14876878 | 0.00531317 | 1.226342 |
| rIP | 0.0554411 | 0.0050401 | 1.07341728 |
| rPT | 0.05313592 | 0.00483054 | 0.9560589 |
| rI | 0.05312278 | 0.00482934 | 0.95539036 |
| lRAC | 0.05111165 | 0.00464651 | 0.8530022 |
| rPOr | 0.05043495 | 0.004585 | 0.81855053 |
| lPT | 0.04928521 | 0.00448047 | 0.76001677 |
| lSM | 0.04908545 | 0.00446231 | 0.74984663 |
| Vocabulary | 0.12007503 | 0.00428839 | 0.65244852 |
| Attention | 0.11938511 | 0.00426375 | 0.63864972 |
| rRMF | 0.0461338 | 0.00419398 | 0.59957635 |
| rSP | 0.03557119 | 0.00323374 | 0.06182599 |
| rSM | 0.02931132 | 0.00266467 | -0.2568687 |
| rCAC | 0.02929821 | 0.00266347 | -0.2575361 |
| lPOr | 0.02785309 | 0.0025321 | -0.331108 |
| lCAC | 0.02722021 | 0.00247456 | -0.3633284 |
| lPOp | 0.02717348 | 0.00247032 | -0.3657076 |
| rPOp | 0.02573646 | 0.00233968 | -0.4388673 |
| Anger | 0.07159169 | 0.00230941 | -0.4558183 |
| lRMF | 0.02273539 | 0.00206685 | -0.5916536 |
| lCMF | 0.02120297 | 0.00192754 | -0.6696706 |
| Executive | 0.05077801 | 0.0018135 | -0.7335361 |
| rRAC | 0.01986409 | 0.00180583 | -0.7378339 |
| rSF | 0.01940129 | 0.00176375 | -0.7613955 |
| Fear | 0.05217904 | 0.00168319 | -0.8065097 |
| Sadness | 0.04351265 | 0.00140363 | -0.9630689 |
| lSP | 0.01329788 | 0.0012089 | -1.0721242 |
| Speed | 0.02609317 | 0.0009319 | -1.2272487 |
| lIP | 0 | 0 | -1.7491289 |
| lSF | 0 | 0 | -1.7491289 |
| rCMF | 0 | 0 | -1.7491289 |
